# Thermotherapy has Sexually Dimorphic Responses in APP/PS1 Mice

**DOI:** 10.1101/2024.03.26.586836

**Authors:** Samuel A. McFadden, Mackenzie R. Peck, Lindsey N. Sime, MaKayla F. Cox, Erol D. Ikiz, Caleigh A. Findley, Kathleen Quinn, Yimin Fang, Andrzej Bartke, Erin R. Hascup, Kevin N. Hascup

## Abstract

A thermoregulatory decline occurs with age due to changes in muscle mass, vasoconstriction, and metabolism that lowers core body temperature (Tc). Although lower Tc is a biomarker of successful aging, we have previously shown this worsens cognitive performance in the APP/PS1 mouse model of Alzheimer’s disease (AD) [1]. We hypothesized that elevating Tc with thermotherapy would improve metabolism and cognition in APP/PS1 mice. From 6-12 months of age, male and female APP/PS1 and C57BL/6 mice were chronically housed at 23 or 30°C. At 12 months of age, mice were assayed for insulin sensitivity, glucose tolerance, and spatial cognition. Plasma, hippocampal, and peripheral (adipose, hepatic, and skeletal muscle) samples were procured postmortem and tissue-specific markers of amyloid accumulation, metabolism, and inflammation were assayed. Chronic 30°C exposure increased Tc in all groups except female APP/PS1 mice. All mice receiving thermotherapy had either improved glucose tolerance or insulin sensitivity, but the underlying processes responsible for these effects varied across sexes. In males, glucose regulation was influenced predominantly by hormonal signaling in plasma and skeletal muscle glucose transporter 4 expression, whereas in females, this was modulated at the tissue level. Thermotherapy improved spatial navigation in male C57BL/6 and APP/PS1 mice, with the later attributed to reduced hippocampal soluble amyloid-β (Aβ)_42_. Female APP/PS1 mice exhibited worse spatial memory recall after chronic thermotherapy. Together, the data highlights the metabolic benefits of passive thermotherapy, but future studies are needed to determine therapeutic benefits for those with AD.

## Introduction

Alzheimer’s disease (AD) is characterized by a progressive deterioration in new learning and memory. This neurodegenerative disorder has multiple etiologies [2], with aging identified as the primary risk factor. Factors associated with normal aging may hasten disease progression. For example, basal metabolic rate declines with age due to deficits in muscle mass, vasoconstriction, glucose metabolism, insulin signaling and thermoregulation [3]. All of which leads to a reduction of core body temperature (Tc) [4].

Low Tc is a biomarker of longevity [5,6] and exposure to hypothermic environmental temperatures triggers adaptive metabolic enhancements. However, the advantageous aspects of lower Tc may adversely affect cognition. Mild hypothermia impairs cognitive performance in rats [7,8], nonhuman primates [9], and humans [10]. Alertness and cognition are strongly correlated with Tc whereby maximal mental performance is observed at higher temperatures [11]. Additionally, working, short-term, and long-term memory performance is decreased in humans during endogenous periods of lower Tc [11–13]. *In vitro* studies have demonstrated that amyloid fibril formation and tau hyperphosphorylation, both pathological hallmarks associated with AD, are accelerated at lower temperatures [14,15]. This potentially indicates a dualistic relationship between the mechanisms facilitating successful physiological aging and those contributing to pathological disease progression.

In our prior study, we demonstrated that APP/PS1 mice subjected to chronic 16°C conditions exhibited improved insulin sensitivity. However, spatial learning and memory recall remained impaired, with females experiencing more pronounced deficits compared to littermates housed at ambient (23°C) temperatures [1]. We also observed increased hippocampal plaque burden in male APP/PS1 mice, similar to observations in 3xTg mice when exposed to hypothermic environmental temperature [16]. Alternatively, increasing Tc may be an effective strategy to treat AD. Passive thermotherapy positively modulates health benefits across physical, cardiovascular, and metabolic disorders [17] that are known AD risk factors [2].

We hypothesized that elevating Tc would enhance metabolism, thereby resulting in improved learning and memory in the amyloidogenic APP/PS1 AD mouse model. Double transgenic APP/PS1 mice express a chimeric mouse/human amyloid precursor protein (APP, Swe695) and a mutant human presenilin (PS)1 lacking exon 9 (ΔE9). These mutations overexpress the amyloid precursor protein with preferential cleavage of amyloid-β (Aβ)_42_ isoforms [18]. Amyloid accumulation and subtle cognitive impairments are observed at 6 months of age and become prominent by 12 months [19]. We have previously shown APP/PS1 mice develop impairments in insulin sensitivity and glucose homeostasis that contribute to their cognitive deficits [20]. Starting at 6 months of age APP/PS1 and C57BL/6 littermate control mice were chronically housed in a 30°C controlled environmental chamber for 6 months. Translationally, this time frame corresponds to conversion from MCI to AD, allowing us to determine the effects of mild hyperthermia on metabolism and cognitive function during disease progression.

## Results

### Core Body Temperature

Tc was determined in a cohort of 11-12 month old mice maintained at ambient temperature since birth. Mice were then chronically exposed to thermotherapy (30°C) for one month before Tc was measured. Thermotherapy increased Tc in all groups of mice except female APP/PS1 (Figure 1). The descriptive statistics for all bar graphs are shown in Table 1.

**Figure 1:**
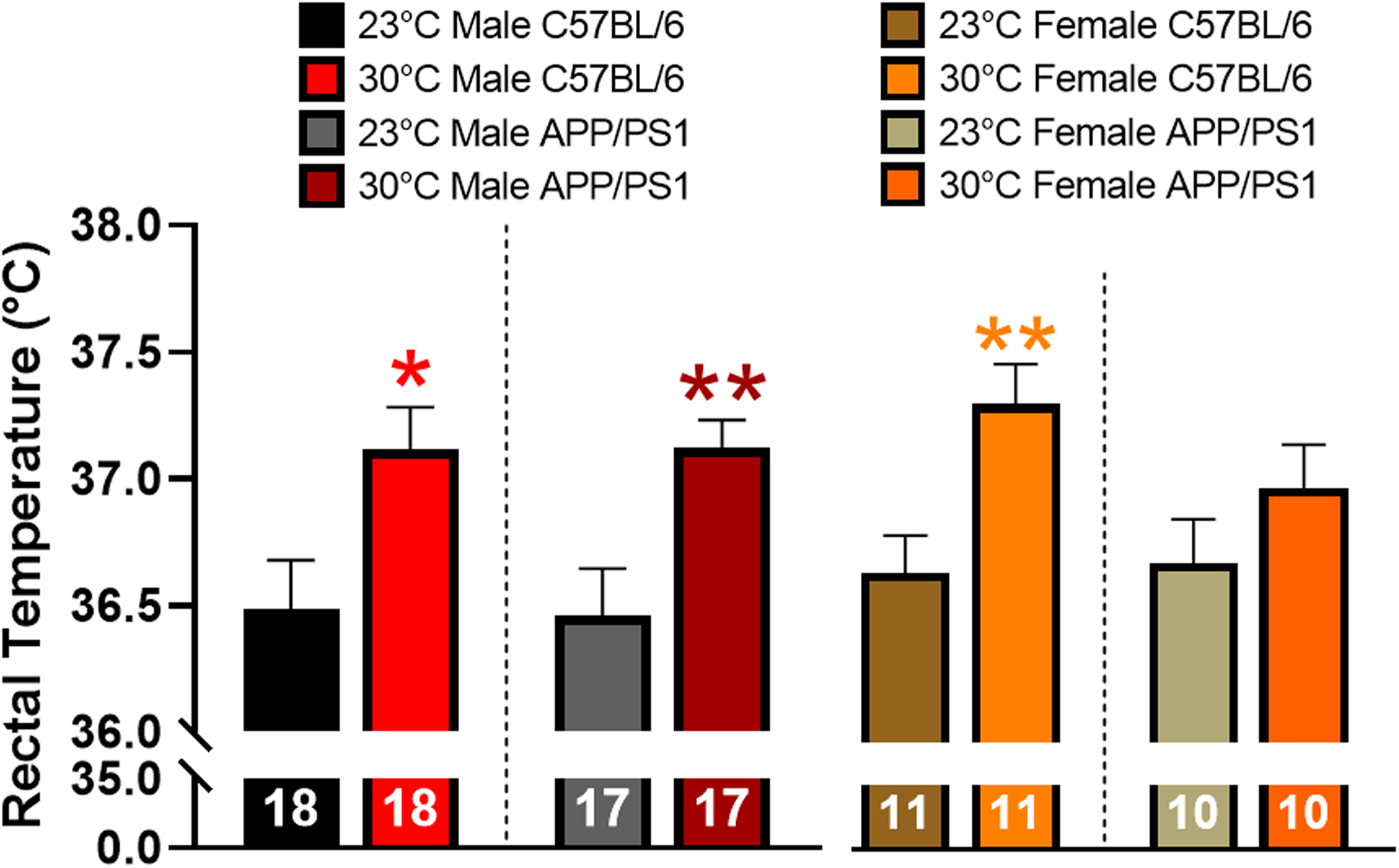
Effects of thermotherapy on Tc. Rectal temperature was determined before (23°C) and after one month of thermotherapy treatment (30°C). The number of animals is inset on each bar graph. A two-tailed t-test was used to determine changes in Tc within a genotype. *p<0.05, **p<0.01.

**Table 1:**
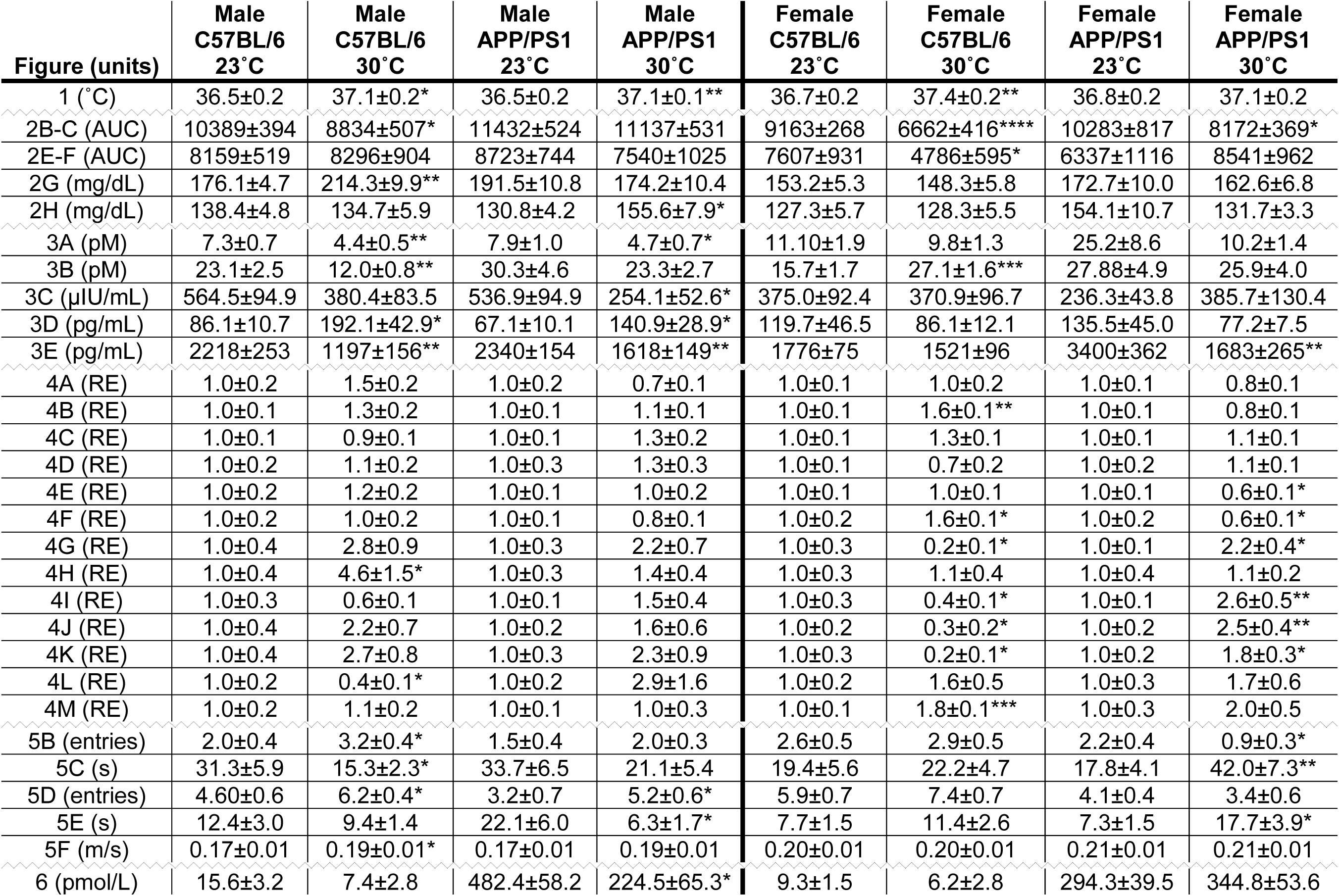
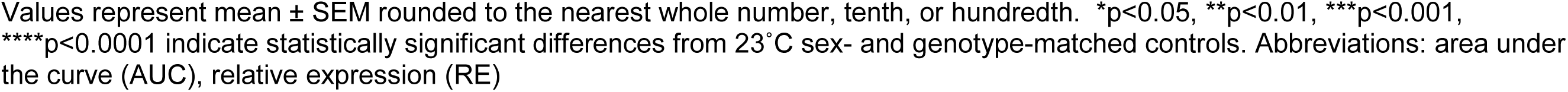
Descriptive statistics for all bar graphs.

| Figure (units) | Male<br>C57BL/6<br>23°C | Male<br>C57BL/6<br>30°C | Male<br>APP/PS1<br>23°C | Male<br>APP/PS1<br>30°C | Female<br>C57BL/6<br>23°C | Female<br>C57BL/6<br>30°C | Female<br>APP/PS1<br>23°C | Female<br>APP/PS1<br>30°C |
| --- | --- | --- | --- | --- | --- | --- | --- | --- |
| 1 (°C) | 36.5±0.2 | 37.1±0.2* | 36.5±0.2 | 37.1±0.1** | 36.7±0.2 | 37.4±0.2** | 36.8±0.2 | 37.1±0.2 |
| 2B-C (AUC) | 10389±394 | 8834±507* | 11432±524 | 11137±531 | 9163±268 | 6662±416**** | 10283±817 | 8172±369* |
| 2E-F (AUC) | 8159±519 | 8296±904 | 8723±744 | 7540±1025 | 7607±931 | 4786±595* | 6337±1116 | 8541±962 |
| 2G (mg/dL) | 176.1±4.7 | 214.3±9.9** | 191.5±10.8 | 174.2±10.4 | 153.2±5.3 | 148.3±5.8 | 172.7±10.0 | 162.6±6.8 |
| 2H (mg/dL) | 138.4±4.8 | 134.7±5.9 | 130.8±4.2 | 155.6±7.9* | 127.3±5.7 | 128.3±5.5 | 154.1±10.7 | 131.7±3.3 |
| 3A (pM) | 7.3±0.7 | 4.4±0.5** | 7.9±1.0 | 4.7±0.7* | 11.10±1.9 | 9.8±1.3 | 25.2±8.6 | 10.2±1.4 |
| 3B (pM) | 23.1±2.5 | 12.0±0.8** | 30.3±4.6 | 23.3±2.7 | 15.7±1.7 | 27.1±1.6*** | 27.88±4.9 | 25.9±4.0 |
| 3C (μIU/mL) | 564.5±94.9 | 380.4±83.5 | 536.9±94.9 | 254.1±52.6* | 375.0±92.4 | 370.9±96.7 | 236.3±43.8 | 385.7±130.4 |
| 3D (pg/mL) | 86.1±10.7 | 192.1±42.9* | 67.1±10.1 | 140.9±28.9* | 119.7±46.5 | 86.1±12.1 | 135.5±45.0 | 77.2±7.5 |
| 3E (pg/mL) | 2218±253 | 1197±156** | 2340±154 | 1618±149** | 1776±75 | 1521±96 | 3400±362 | 1683±265** |
| 4A (RE) | 1.0±0.2 | 1.5±0.2 | 1.0±0.2 | 0.7±0.1 | 1.0±0.1 | 1.0±0.2 | 1.0±0.1 | 0.8±0.1 |
| 4B (RE) | 1.0±0.1 | 1.3±0.2 | 1.0±0.1 | 1.1±0.1 | 1.0±0.1 | 1.6±0.1** | 1.0±0.1 | 0.8±0.1 |
| 4C (RE) | 1.0±0.1 | 0.9±0.1 | 1.0±0.1 | 1.3±0.2 | 1.0±0.1 | 1.3±0.1 | 1.0±0.1 | 1.1±0.1 |
| 4D (RE) | 1.0±0.2 | 1.1±0.2 | 1.0±0.3 | 1.3±0.3 | 1.0±0.1 | 0.7±0.2 | 1.0±0.2 | 1.1±0.1 |
| 4E (RE) | 1.0±0.2 | 1.2±0.2 | 1.0±0.1 | 1.0±0.2 | 1.0±0.1 | 1.0±0.1 | 1.0±0.1 | 0.6±0.1* |
| 4F (RE) | 1.0±0.2 | 1.0±0.2 | 1.0±0.1 | 0.8±0.1 | 1.0±0.2 | 1.6±0.1* | 1.0±0.2 | 0.6±0.1* |
| 4G (RE) | 1.0±0.4 | 2.8±0.9 | 1.0±0.3 | 2.2±0.7 | 1.0±0.3 | 0.2±0.1* | 1.0±0.1 | 2.2±0.4* |
| 4H (RE) | 1.0±0.4 | 4.6±1.5* | 1.0±0.3 | 1.4±0.4 | 1.0±0.3 | 1.1±0.4 | 1.0±0.4 | 1.1±0.2 |
| 4I (RE) | 1.0±0.3 | 0.6±0.1 | 1.0±0.1 | 1.5±0.4 | 1.0±0.3 | 0.4±0.1* | 1.0±0.1 | 2.6±0.5** |
| 4J (RE) | 1.0±0.4 | 2.2±0.7 | 1.0±0.2 | 1.6±0.6 | 1.0±0.2 | 0.3±0.2* | 1.0±0.2 | 2.5±0.4** |
| 4K (RE) | 1.0±0.4 | 2.7±0.8 | 1.0±0.3 | 2.3±0.9 | 1.0±0.3 | 0.2±0.1* | 1.0±0.2 | 1.8±0.3* |
| 4L (RE) | 1.0±0.2 | 0.4±0.1* | 1.0±0.2 | 2.9±1.6 | 1.0±0.2 | 1.6±0.5 | 1.0±0.3 | 1.7±0.6 |
| 4M (RE) | 1.0±0.2 | 1.1±0.2 | 1.0±0.1 | 1.0±0.3 | 1.0±0.1 | 1.8±0.1*** | 1.0±0.3 | 2.0±0.5 |
| 5B (entries) | 2.0±0.4 | 3.2±0.4* | 1.5±0.4 | 2.0±0.3 | 2.6±0.5 | 2.9±0.5 | 2.2±0.4 | 0.9±0.3* |
| 5C (s) | 31.3±5.9 | 15.3±2.3* | 33.7±6.5 | 21.1±5.4 | 19.4±5.6 | 22.2±4.7 | 17.8±4.1 | 42.0±7.3** |
| 5D (entries) | 4.60±0.6 | 6.2±0.4* | 3.2±0.7 | 5.2±0.6* | 5.9±0.7 | 7.4±0.7 | 4.1±0.4 | 3.4±0.6 |
| 5E (s) | 12.4±3.0 | 9.4±1.4 | 22.1±6.0 | 6.3±1.7* | 7.7±1.5 | 11.4±2.6 | 7.3±1.5 | 17.7±3.9* |
| 5F (m/s) | 0.17±0.01 | 0.19±0.01* | 0.17±0.01 | 0.19±0.01 | 0.20±0.01 | 0.20±0.01 | 0.21±0.01 | 0.21±0.01 |
| 6 (pmol/L) | 15.6±3.2 | 7.4±2.8 | 482.4±58.2 | 224.5±65.3* | 9.3±1.5 | 6.2±2.8 | 294.3±39.5 | 344.8±53.6 |

Values represent mean ± SEM rounded to the nearest whole number, tenth, or hundredth. \* $p < 0.05$ , \*\* $p < 0.01$ , \*\*\* $p < 0.001$ , \*\*\*\* $p < 0.0001$ indicate statistically significant differences from 23°C sex- and genotype-matched controls. Abbreviations: area under the curve (AUC), relative expression (RE)

### Thermotherapeutic effects on Blood Glucose

Repeated exposure (3-6 times per week) to thermotherapy for 30-60 minutes positively modulates insulin sensitivity and glucose tolerance in people with metabolic dysfunction [21,22]. Previous research shows APP/PS1 mice have reduced insulin sensitivity and glucose tolerance compared with age-matched littermate controls [1,20,23]. To determine thermotherapeutic effects on blood glucose regulation, we performed an insulin and glucose tolerance tests (ITT and GTT, respectively) after six months of chronic thermotherapy treatment. During the ITT, an intraperitoneal (ip) injection of insulin decreased blood glucose levels to a greater extent in all groups receiving thermotherapy, except for male APP/PS1, when compared with genotype and sex-matched ambient temperature controls (Figure 2A). The ITT area under the curve (AUC) provides an overall indication of insulin sensitivity. Thermotherapy improved insulin sensitivity in all groups apart from male APP/PS1 mice (Figure 2B-C). The GTT provides an indication of the endogenous uptake of blood sugar into tissue for energy utilization or storage. Thermotherapy exposed male APP/PS1 and female C57BL/6 mice receiving thermotherapy had lower blood glucose levels 15 minutes after an ip injection of glucose than their corresponding ambient temperature controls (Figure 2D). No time point differences were observed in male C57BL/6 or female APP/PS1 mice. The GTT AUC provides an indication of overall glucose tolerance which was only improved in female C57BL/6 mice receiving thermotherapy (Figure 2E-F). Thermotherapy worsened fed blood glucose in male C57BL/6 (Figure 2G) and fasting blood glucose in male APP/PS1 mice (Figure 2H) despite their improved insulin sensitivity and glucose tolerance, respectively.

**Figure 2:**
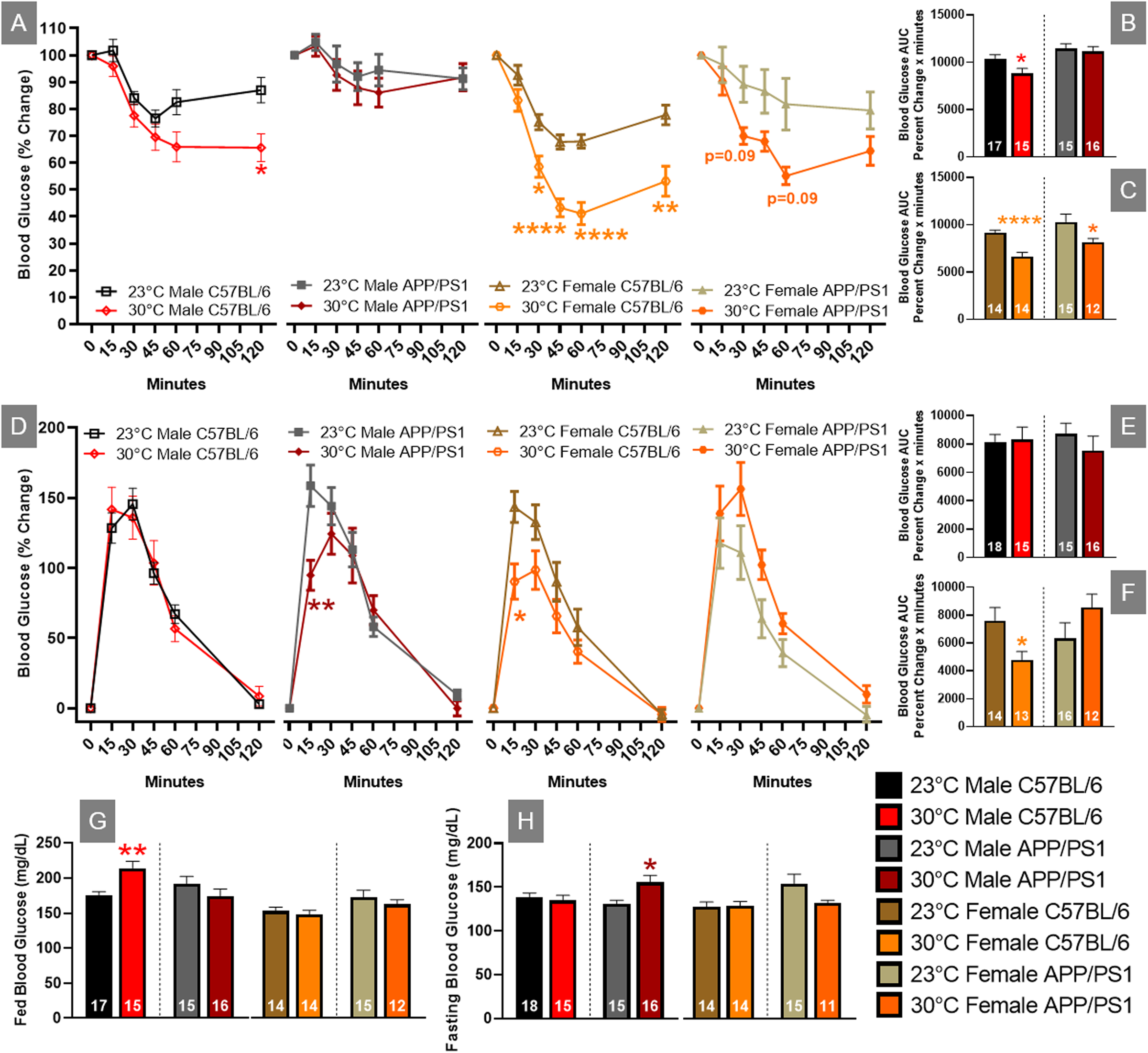
Blood glucose changes after thermotherapy. Percent change of blood glucose levels after an ip injection of 1 IU / kg bw of insulin at t=0 minutes (A). The AUC was calculated for the duration of the ITT and compared temperature effects in male (B) and female (C) mice. Percent change of blood glucose levels after an ip injection of 2 g / kg bw of glucose at t=0 minutes (D). The AUC was calculated for both male (E) and female (F) mice. Fed (G) and fasting (H) blood glucose levels were determined prior to ip injection of insulin and glucose, respectively. The number of animals is inset on each bar graph. A two-way ANOVA factor analysis with Sidak’s post hoc was used to determine significant blood glucose changes due to thermotherapy across time intervals. A two-tailed t-test was used to determine blood glucose changes within a genotype and sex. *p<0.05, **p<0.01, ****p<0.0001.

### Effects of glucose regulating plasma peptides after chronic thermotherapy

Circulating levels of plasma peptides and hormones that modulate blood glucose levels were measured by multiplex assays. Glucagon release from the pancreas promotes liver glycogenolysis thereby increasing circulating blood glucose levels. Chronic thermotherapy reduced plasma glucagon levels in all groups except female C57BL/6 mice (Figure 3A). Glucagon-like peptide 1 (GLP1) release from the gastrointestinal tract reduces circulating glucose levels by stimulating insulin release, suppressing glucagon secretion, and promoting satiety. Thermotherapeutic differences in plasma GLP1 levels were only observed in C57BL/6 mice with decreased levels in males and an increase observed in females (Figure 3B). Despite the changes observed in GLP1, thermotherapy did not affect plasma insulin in C57BL/6 mice, but a decrease was observed in male APP/PS1 mice (Figure 3C). Fibroblast growth factor 21 (FGF21) regulates metabolism and is important for thermogenic recruitment of white adipose tissue (WAT). Plasma FGF21 was elevated in male C57BL/6 and APP/PS1 mice housed at 30°C, but no differences were observed in females receiving thermotherapy (Figure 3D). B-cell activating factor (BAFF) is a member of the tumor necrosis factor (TNF) ligand family that regulates adipose tissue inflammation and impairment of insulin-receptor signaling [24]. Chronic thermotherapy treatment reduced plasma BAFF in both genotypes of male mice as well as female APP/PS1 mice, similar to plasma glucagon observations (Figure 3E). Taken together, these results show that six months of passive thermotherapy altered the plasma profile of peptides and hormones in a manner consistent with improved insulin sensitivity and glucose tolerance in a sex-dependent manner.

**Figure 3:**
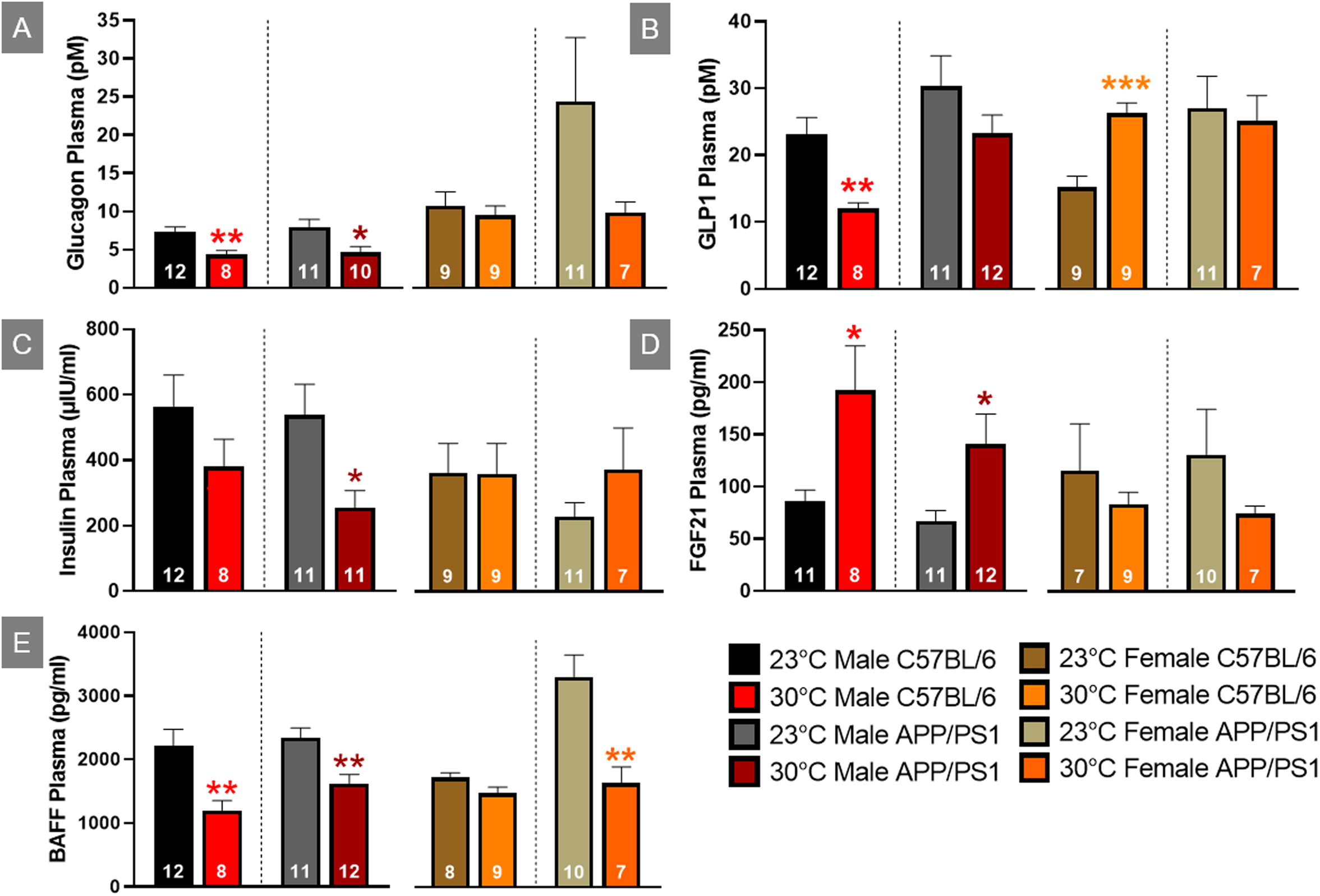
Plasma concentrations of glucose regulating hormones. Plasma expression levels of glucagon (A), glucagon-like peptide 1 (GLP1; B), insulin (C), fibroblast growth factor 21 (FGF21; E), and B-cell activating factor (BAFF; E) as detected by multiplex assay. A two-tailed t-test was used to determine plasma concentration changes within a genotype and sex. *p<0.05, **p<0.01, ***p<0.001.

### Hepatic glucose metabolizing genes

The liver plays a key role in maintaining an easily accessible supply of energy, primarily in the form of glucose and glycogen. Insulin receptor (InsR) activation leads to downstream signal transduction through phosphatidylinositol 3-kinase (PI3K) / Akt that induces translocation of glucose transporters to the cell surface, thereby facilitating glucose uptake. Once in the liver, glucose undergoes phosphorylation by glucokinase (Gck) to form glucose-6-phosphate, which can be utilized in either glycolysis or glycogenesis. Conversely, glucose-6-phosphatase (G6PC) catalyzes the hydrolysis of glucose-6-phosphate back to glucose. Chronic thermotherapy affected genes associated with liver glucose uptake in female C57BL/6 mice. InsR and Akt expression were similar, but PI3K relative gene expression was elevated in after thermotherapy treatment (Figure 4A-C). This increased expression accounts for the improved insulin sensitivity seen in the female C57BL/6 mice as previously discussed (Figure 2A, C). No difference in Glut2 expression were observed (Figure 4D), but GCK and G6PC were both decreased in female APP/PS1 mice receiving thermotherapy treatment (Figure 4E-F). Conversely, thermotherapy increased G6PC expression in female C57BL/6 mice. This variation in G6PC between female C57BL/6 and APP/PS1 mice likely contributes to their divergent responses during the GTT. Following thermotherapy treatment, glucose tolerance was enhanced in C57BL/6, but diminished in APP/PS1 mice.

**Figure 4:**
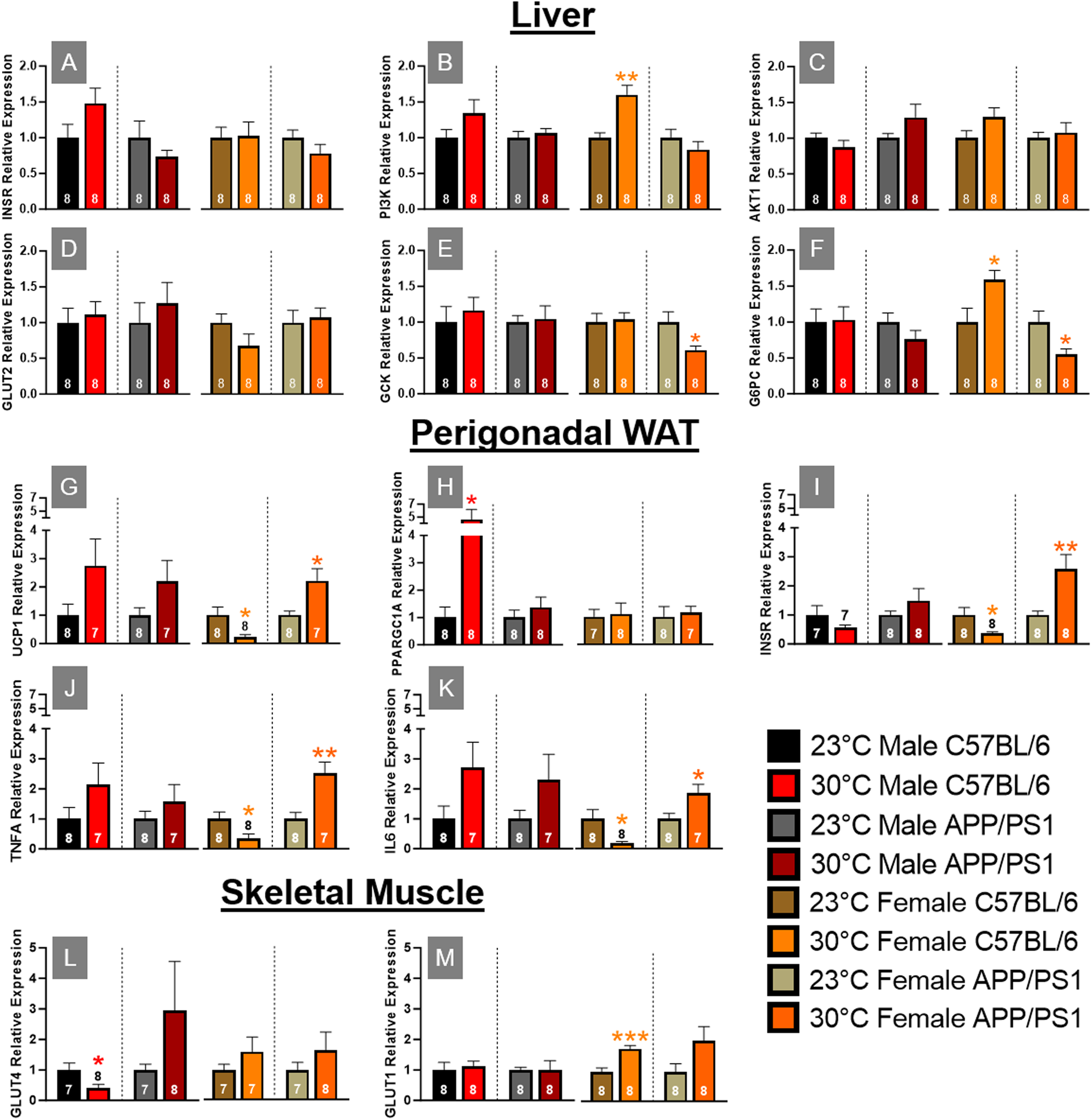
Hepatic, perigonadal white adipose tissue, and skeletal muscle mRNA expression. Hepatic mRNA expression of insulin receptor (INSR; A), phosphatidylinositol 3-kinase (PI3K; B), AKT1 (C), glucose transporter 2 (GLUT2; D), glucokinase (GCK; E), and glucose 6-phosphatase (G6PC; F) relative to B2M. Perigonadal WAT mRNA expression changes of uncoupling protein 1 (UCP1; G), PGC-1α (PPARGC1A; H), INSR (I), tumor necrosis factor α (TNFA; J), and interleukin 6 (IL6; K) relative to B2M. GLUT4 (L) and 1 (M) mRNA expression relative to GAPDH in skeletal muscle. A two-tailed t-test was used to determine mRNA expression fold changes within a genotype and sex. The number of animals is either inset or above the error bars on each bar graph. *p<0.05, **p<0.01, ***p<0.001.

### Perigonadal white adipose tissue mRNA expression

Visceral adipose tissue is hormonally active and regulates systemic metabolism by acting as an energy reservoir and through secretion of adipokines and cytokines. Excessive accumulation of this adipose depot causes insulin insensitivity and release of proinflammatory cytokines resulting in metabolic dysregulation. Localized thermotherapy has been shown to induce browning of WAT through increased expression of uncoupling protein (UCP) 1 [25]. UCP1 is present in brown and beige adipose tissue and is responsible for dissipating the mitochondrial electron transport gradient to generate heat instead of ATP synthesis. Chronic thermotherapy increased perigonadal (pg) WAT UCP1 expression in all groups except female C57BL/6 mice, where a decrease was observed (Figure 4G). Peroxisome proliferator-activated receptor-gamma coactivator (PGC)-1α regulates expression of UCP1 along with FGF21 signaling. PGC-1α expression was upregulated in male C57BL/6 mice after thermotherapy treatment (Figure 4H). This increased gene expression coupled with FGF21 plasma concentrations (Figure 3D) indicates that thermotherapy increased gene expression pathways implicated in the browning of adipose tissue. Thermotherapy modulated InsR expression in female mice, but in opposing manners, with reduced expression in C57BL/6 and an increase observed in APP/PS1 mice (Figure 4I). This would account for the improved insulin sensitivity in female APP/PS1, but not sex-matched C57BL/6 mice. Proinflammatory cytokines systemically impair glucose homeostasis. Similar to InsR expression, TNFα and Interleukin (IL)-6 mRNA levels were divergent in female mice after thermotherapy treatment. In females, expression of both proinflammatory cytokines were decreased in pgWAT of C57BL/6, but increased in APP/PS1 mice (Figure 4J-K). The increased release of proinflammatory cytokines from pgWAT in female APP/PS1 likely contributes to the worsening of glucose tolerance discussed previously (Figure 2D,F).

### Skeletal Muscle Glucose Transporter Expression

Skeletal muscle is the predominant tissue responsible for blood glucose homeostasis mediated through insulin-dependent (Glut4) and independent (Glut1) mechanisms. Under basal conditions, Glut1 is located on the plasma membrane, whereas Glut4 resides intracellularly. In response to insulin or exercise, Glut4 is translocated to the plasma membrane and is crucial for systemic glucose homeostasis. Glut1 can also enhance basal glucose uptake in skeletal muscle [26]. Thermotherapy showed opposing effects on Glut4 expression in male mice with decreases in C57BL/6 but increases in APP/PS1 mice (Figure 4L), while Glut1 expression was similar in both genotypes (Figure 4M). In male APP/PS1 mice, Glut4 expression levels were elevated (although not significantly) that could contribute to improving their glucose tolerance (Figure 2D). Whereas in male C57BL/6 mice, decreased Glut4 expression after thermotherapy does not affect glucose tolerance despite improvements in insulin signaling. In females, thermotherapy did not alter Glut4 but increased expression of Glut1 in C57BL/6 mice (Figure 4L-M). This enhanced Glut1 expression in female C57BL/6 would account for the improvement in glucose tolerance discussed earlier (Figure 2D), particularly at the earliest time point.

### Thermotherapy alters spatial learning and memory in a sexually dimorphic manner

Spatial learning and memory recall is impaired in APP/PS1 mice by 12 months of age [1]. To determine if thermotherapy ameliorates these cognitive deficits, we tested mice using the Morris water maze (MWM) spatial navigation paradigm. During the training sessions, platform latency was decreased in male C57BL/6 mice exposed to 30°C (Figure 5A). All other groups had a similar learning profile. During the probe challenge, the number of platform entries (Figure 5B) were increased while latency to first platform entry (Figure 5C) was decreased in male C57BL/6 mice after thermotherapy. Female APP/PS1 mice exhibited a decrease in the number of platform entries and an increase in platform latency for first entry (Figure 5B-C). When examining an area slightly larger than the former location of the hidden escape platform, the number of annulus 40 entries (Figure 5D) were increased in both male C57BL/6 and APP/PS1 mice receiving thermotherapy. This also reduced the latency to first annulus 40 entry in male APP/PS1 mice (Figure 5E), but an increased latency in female APP/PS1 mice was observed. Thermotherapy increased the swimming speed in male C57BL/6 mice only (Figure 5F) that may be indicative of skeletal muscle enhancements due to the warmer environment [27]. This faster swimming along with an improved learning curve in male C57BL/6 mice accounts for their improved memory recall. However, in AD mice, these performance issues were not observed. Thermotherapy has sexually dimorphic cognitive effects regardless of AD genotype.

**Figure 5:**
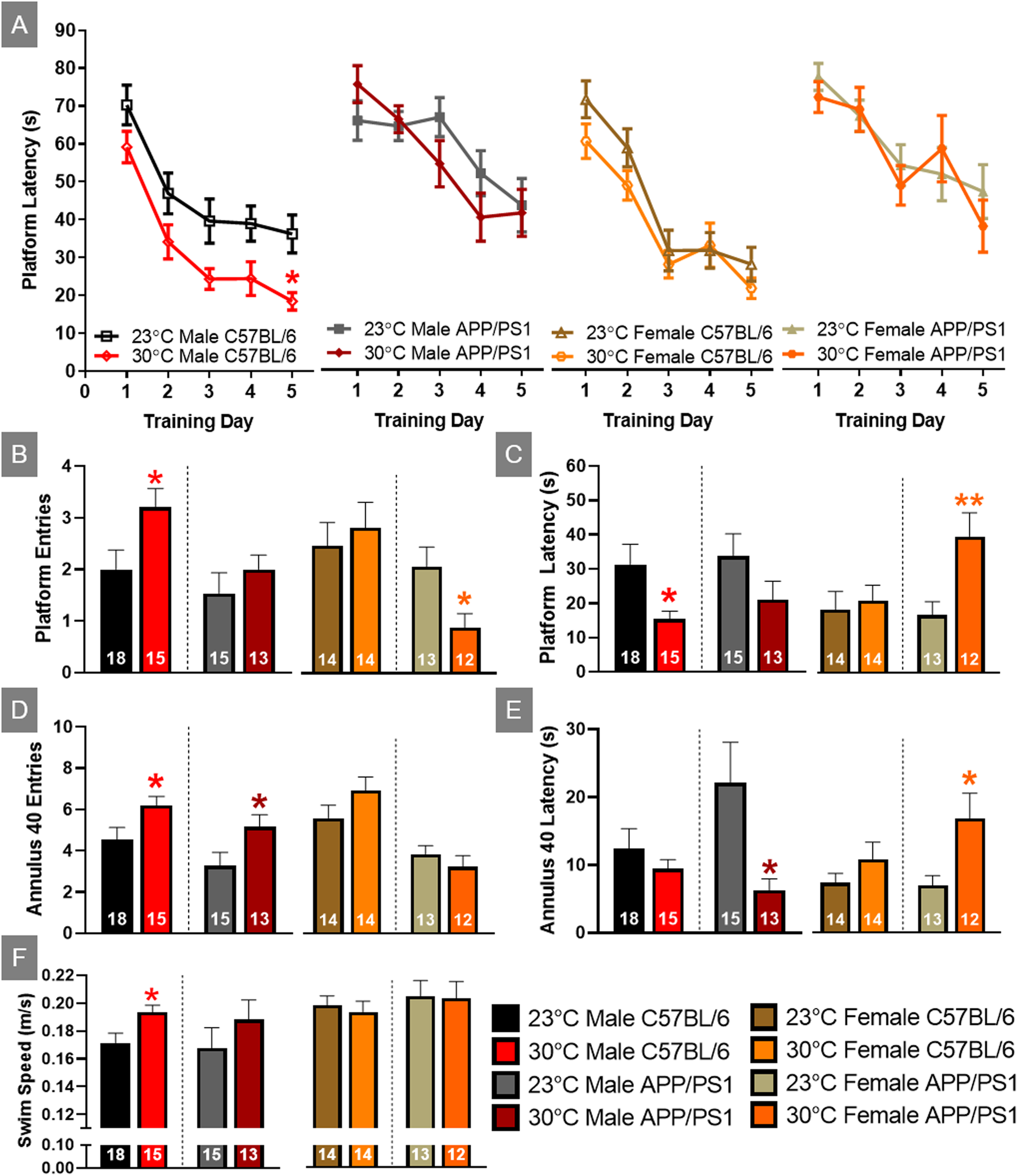
Sexually dimorphic spatial navigation responses to thermotherapy in APP/PS1 mice. Average latency to the platform during the five MWM training sessions (A). A two-way ANOVA with Sidak’s post hoc was used to determine significant time differences due to thermotherapy across training days. The number of platform entries (B), latency to first platform entry (C), number of annulus 40 entries (D), latency to first annulus 40 entry (E), and swimming speed (F) during the delayed MWM probe challenge. A two-tailed t-test was used to determine thermotherapy effects on probe challenge parameters within a sex and genotype. The number of animals is inset on each bar graph. *p<0.05, **p<0.01.

### Thermotherapy reduces hippocampal soluble Aβ_42_ concentration in male APP/PS1 mice

Soluble Aβ_42_ is considered to be a neurotoxic species and its aggregation of monomers, oligomers, and protofibrils are associated with cognitive decline in AD [28]. Transgenic APP/PS1 mice overexpress APP and PS1 resulting in preferential cleavage of the 42 base pair Aβ peptide. A soluble Aβ_42_ specific ELISA was used to measure hippocampal concentrations in APP/PS1 and C57BL/6 mice, with the latter serving as a negative control (Figure 6). Thermotherapy reduced the hippocampal Aβ_42_ concentration in APP/PS1 male mice, but no effects were observed in females of either genotype. This reduction of Aβ_42_ in male APP/PS1 mice likely contributes to the improved spatial memory recall.

**Figure 6:**
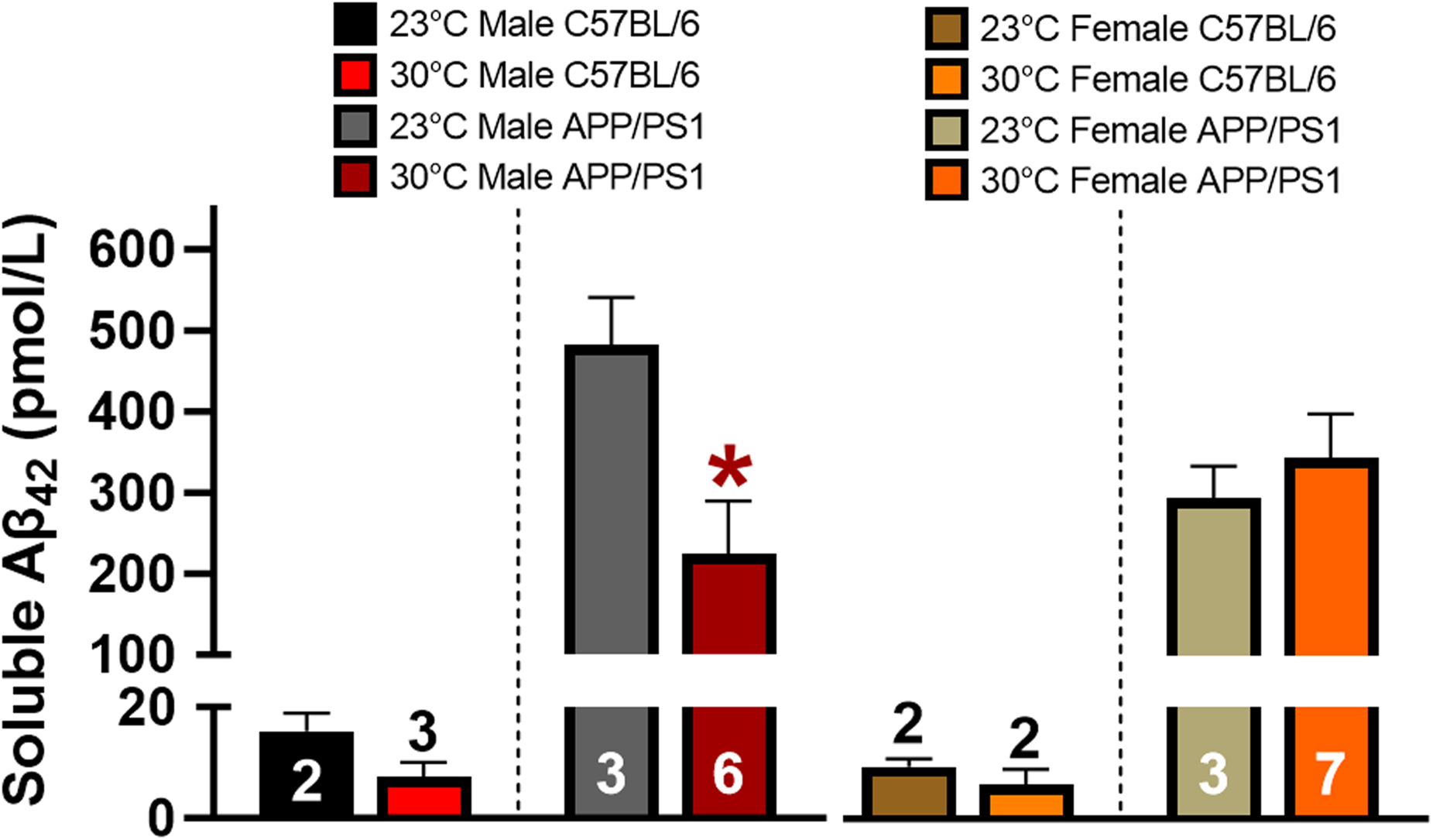
Thermotherapy decreases hippocampal soluble Aβ_42_ in male mice. Average concentration of hippocampal soluble Aβ_42_ was determined by ELISA in male and female mice. C57BL/6 mice were used as a negative control as denoted by the segmented y-axis. A two-tailed t test was used to determine thermotherapy effects within a sex and genotype. The number of animals is either inset or above the error bars on each bar graph. *p<0.05.

## Discussion

The age of the mice for this study was chosen based on the pathology and cognitive aspects of disease progression. Amyloid accumulation and subtle cognitive impairments are observed at 6 months of age and rapidly progress to 12 months in APP/PS1 mice. This time frame translationally corresponds to mild cognitive impairment which is the opportune window to initiate disease-modifying treatment to prevent conversion to AD. Various biological and environmental factors contribute to the onset and progression of AD, leading to the development of cognitive deficits as well as metabolic and physical decline. To improve the overall quality of life for individuals with AD, effective treatments should target a range of physiological changes beyond cognitive impairments. Passive thermotherapy positively modulates multiple physiological parameters and represents a nonpharmacological approach for potential disease modifying treatment. Polypharmacy is common among aging populations, and nonpharmacological interventions help mitigate adverse drug reactions.

Despite passive thermotherapy increasing Tc in both sexes of C57BL/6 mice, we observed some notable differences between the sexes. Insulin sensitivity was improved in both sexes, but to a much larger degree in the females. Improved glucose tolerance was only observed in female C57BL/6 mice. Thermotherapy significantly influenced plasma hormone levels in male C57BL/6 mice, while mRNA expression levels were more affected in females. In male mice, plasma glucagon, GLP1, and BAFF levels decreased, whereas FGF21 levels increased. In female mice, only GLP1 levels showed an increase. The increased expression of hepatic PI3K, combined with the decreased expression of INSR in pgWAT, likely contributed to the improved insulin sensitivity observed in female C57BL/6 mice following thermotherapy treatment. Additionally, we observed reduced expression levels of proinflammatory cytokines in the pgWAT of these female mice. Finally, thermotherapy improved spatial learning and memory only in male C57BL/6 mice, but this may be attributed to improved muscle performance since swimming speed was faster in these mice.

Our present study shows that passive thermotherapy improved either glucose tolerance or insulin sensitivity in all sexes and genotypes tested. The underlying processes responsible for these effects are diverse and sexually dimorphic. In males, glucose regulation was influenced predominantly by plasma hormone signaling and skeletal muscle Glut4 expression, whereas in females, this was modulated at the level of individual tissues. Reduced plasma glucagon levels decrease glycogenolysis, improved insulin sensitivity accompanies lower BAFF concentrations, and skeletal muscle Glut4 expression regulates uptake of circulating blood glucose levels. An increase in plasma FGF21 and elevated gene expression of UCP1 and PGC-1α in pgWAT indicates thermotherapy induced WAT browning that improved metabolism, similar to previous observations [25]. Female mice receiving thermotherapy treatment showed significantly enhanced responses to insulin that were mediated by changes in plasma hormones and pgWAT genes. In female C57BL/6 mice, thermotherapy enhanced insulin signaling and lowered blood glucose levels. This was accompanied by elevated plasma concentrations of GLP1 and hepatic PI3K gene expression. In thermotherapy treated female APP/PS1 mice, decreased BAFF plasma concentrations and increased pgWAT InsR expression would also improve their response to insulin. However, the elevated expression of proinflammatory cytokines in pgWAT after thermotherapy would cause the slower glucose clearance in these mice.

Despite the metabolic improvements observed in both sexes after thermotherapy, cognitive improvements were only observed in male mice. Soluble Aβ_42_, rather than insoluble fibrils and plaques, is considered neurotoxic due to its ability to spread throughout the brain and affect energy metabolism, neurotransmission, and inflammation. This eventually causes synapse and cell loss that is a contributing factor to cognitive decline. Soluble Aβ_42_ was reduced in males of both genotypes, consistent with their improved spatial navigation. Although thermotherapy did not affect soluble Aβ_42_ levels in female APP/PS1 mice, their memory recall was significantly decreased. This could be attributed to the elevated proinflammatory cytokine gene expression observed peripherally. In addition to affecting glucose tolerance, TNFα and IL-6 can readily cross the blood brain barrier and exacerbate the neuroinflammation already caused by plaque formation.

Both passive cooling and thermotherapy have been explored in different models of AD and have shown varying results. Acute and chronic passive cooling negatively affected AD pathology and cognition in the 3xTg [16] and the APP/PS1 [1] models of AD, respectively. In the 3xTg model, repeated bouts of passive cooling reduced tau phosphorylation, but had no effect on amyloid pathology. The cognitive implications of these findings were not determined [29]. Thermotherapy reduced soluble Aβ_42_ [16] and tau phosphorylation [30] while improving cognitive performance in the 3xTg mice, which is similar to the male APP/PS1 mice assayed in the present study. Unlike males, chronic thermotherapy in female APP/PS1 mice worsened cognitive performance similar to findings in Tg2576 mice [31].

In our present study, the APP/PS1 female mice were the only group that did not experience an increase in Tc after thermotherapy and, coincidentally, exhibited poorer performance on the MWM paradigm compared with normothermic controls matched for genotype and sex. Despite Tc modulation by estradiol and progesterone [32], the absence of a response is not solely attributed to sex hormones, as thermotherapy increased Tc in female C57BL/6 mice. In mice, thermoregulatory sex differences have not been fully elucidated, but females prefer warmer environments than males regardless of gonadal factors [32] despite having higher Tc [33]. This suggests the differential responses we observed in the present study were not driven by sex hormones, but rather Aβ_42_ accumulation and aggregation. Further research is needed to determine how this accumulation might affect hypothalamic preoptic area modulation of Tc.

Humans and other mammals are homeothermic, able to maintain a stable Tc through their metabolic activity, regardless of external environmental influences. Most people in developed nations spend a majority of their time residing in temperature controlled environments that are optimized for thermal comfort, but minimize thermogenesis [34]. This coupled with the increasingly sedentary nature of work and life reduces activity-related heat production resulting in thermostasis [35]. Accordingly, exposure to thermal extremes has gained scientific interest for its numerous physiological benefits that could either reduce risk factors associated with AD onset or be a treatment for disease progression [17]. Japanese ofuro and Scandinavian sauna bathing has been used for centuries with the latter associated with reduced risk of dementia and AD [36]. Studies demonstrate that exercise and passive heating have comparable beneficial physiological effects [37]. Hence, passive thermotherapy would be a preferable alternative to exercise, especially in elderly and frail individuals.

A couple of limitations should be considered when interpreting the findings of this study. Protein levels rather than gene expression would provide a more causal mechanistic underpinning to the observed metabolic differences that were reported. In particular, mRNA expression does not infer recruitment of glucose transporters to the plasma surface and this analysis would be a better measure of changes in glucose tolerance. Finally, APP/PS1 females undergo reproductive senescence at earlier ages than C57BL/6 mice which could influence glucose metabolism and cognitive performance. To avoid these cofounds, we analyzed within genotype treatment comparisons.

Our present research adds to a growing body of literature highlighting the benefits of passive thermotherapy to modulate AD progression and cognition in male, but not female mice, despite observing greater metabolic effects in the latter. While these sexually dimorphic responses have not been fully elucidated, further research is needed to improve the translational applications, such as investigating the duration of thermotherapy treatment and determining if similar effects are observed regardless of disease severity or pathological progression. Both of these factors are ongoing research efforts in our laboratory. The metabolic benefits highlighted in the present study also have translatable implications to other research areas such as obesity, diabetes, or metabolic syndrome. Finally, thermotherapy can be applied without causing drug-drug interactions and could replace the health benefits of exercise in frailer individuals.

## Methods

### Animals

Male and female APP/PS1 and littermate control C57BL/6 mice used for this study were bred and maintained in our animal colony and originated from founder C57BL/6J (RRID:IMSR_JAX:000664) and APP/PS1 (RRID:MMRRC_034832-JAX) mice from Jackson Laboratory (Bar Harbor, ME). A 5 mm tail tip was sent to TransnetYX^®^, Inc (Cordova, TN) to confirm genotypes. Mice were group-housed according to sex and genotype on a 12:12 hour light / dark cycle, and laboratory rodent diet (LabDiet, 5001) and water were available *ad libitum*. The *in vivo* assays were performed in the same order (ITT, GTT, MWM) for all mice with a minimum of one week apart to limit effects of stress. One week post cognitive assessment, mice were deeply anesthetized with isoflurane and a cardiac puncture for blood chemistry analysis was performed. Immediately following, mice were euthanized by decapitation. Tissues were extracted and stored at −80°C until processing.

### Chemicals

Unless otherwise noted, all chemicals were prepared and stored according to manufacturer recommendations.

### Thermotherapy

From 6 to 12 months of age, mouse cages were placed into an environmental chamber (Powers Scientific Cat: RIS33SD) maintained at 30 ± 1°C located within the same animal facility room. Mice were only removed from this chamber for cage cleanings. A separate cohort of age- and sex-matched C57BL/6 and APP/PS1 mice were maintained at a standard animal room ambient temperature (23 ± 1°C) and used as a within genotype temperature control.

### Core Body Temperature Measurements

A separate cohort of male and female C57BL/6 and APP/PS1 mice were maintained at standard ambient temperature from birth until 11 months of age. A rodent thermometer with rectal probe (TK8851; BioSebLab) was used to obtain Tc when mice were 11 months old. These mice were then transferred to environmental chambers (Powers Scientific Cat: RIS33SD) maintained at 30°C until 12 months of age (1 month chronic treatment). Mice were transferred to a separate room within our animal facility maintained at 30°C to assess changes in Tc after one month of chronic exposure.

### Intraperitoneal (ip) Insulin Tolerance test (ITT) and Glucose Tolerance Test (GTT)

To determine insulin sensitivity, an initial blood glucose measurement (time = 0) was taken from the tail vein of fed mice and measured using a Presto® glucometer (AgaMatrix, Salem, NH) followed by ip injection of 1 IU / kg bw Humulin® R (Henry Schein, Melville, NY: Cat: 1238578). To determine glucose tolerance, an initial blood glucose measurement was taken (time = 0) from fifteen hour fasted mice followed by an ip injection of 2 g of dextrose / kg bw (Fisher Scientific Cat: D15). Following either injection, blood glucose levels were measured sequentially at 15, 30, 45, 60, and 120 min [20].

### Morris Water Maze (MWM) Training and Probe Challenge

At approximately 12 months of age, mice underwent cognitive assessment using the MWM spatial learning and memory recall paradigm, during which mice are trained to utilize visual cues to repeatedly swim to a static, submerged hidden platform. The MWM paradigm consisted of 5 consecutive learning days with three, 90-sec trials/day and a minimum 20 minute inter-trial interval. During the delayed memory recall, the platform was removed and mice were given a single, 60 second probe challenge. The ANY-maze video tracking system (Stoelting Co., Wood Dale, IL; RRID:SCR_014289) was used to record navigational parameters and data analysis. The three trials for each training day were averaged for each mouse for analysis. Variables extracted from ANY-maze and utilized for data analysis include platform entries and latency, annulus 40 entries and latency, and swimming speed.

### Blood Chemistry

A cardiac puncture was used to collect blood in EDTA coated tubes (Sarstedt Inc. Microvette CB 300) on wet ice until centrifugation at 1500 x g for 10 min at 4°C. The plasma supernatant was collected and stored at −80°C until analysis with a multiplex assay kit (Meso Scale Discovery) according to the manufacturer’s recommended protocols.

### RT-PCR

RNA was extracted from tissue and quantified using a NanoDrop One spectrophotometer (Thermo Fisher Scientific) according to our previously published protocols [1]. cDNA was synthesized using candidate primers (Integrated DNA Technologies; Table 2) and an iScript cDNA Synthesis Kit (Bio-Rad). Relative mRNA expression was analyzed by quantitative RT-PCR using the QuantStudio PCR System (Applied Biosystems) and SYBR Green MasterMix (Bio-Rad). Beta-2-microglobulin (B2M) was used as the internal housekeeping gene for pgWAT and liver while GAPDH was used for skeletal muscle.

**Table 2:**
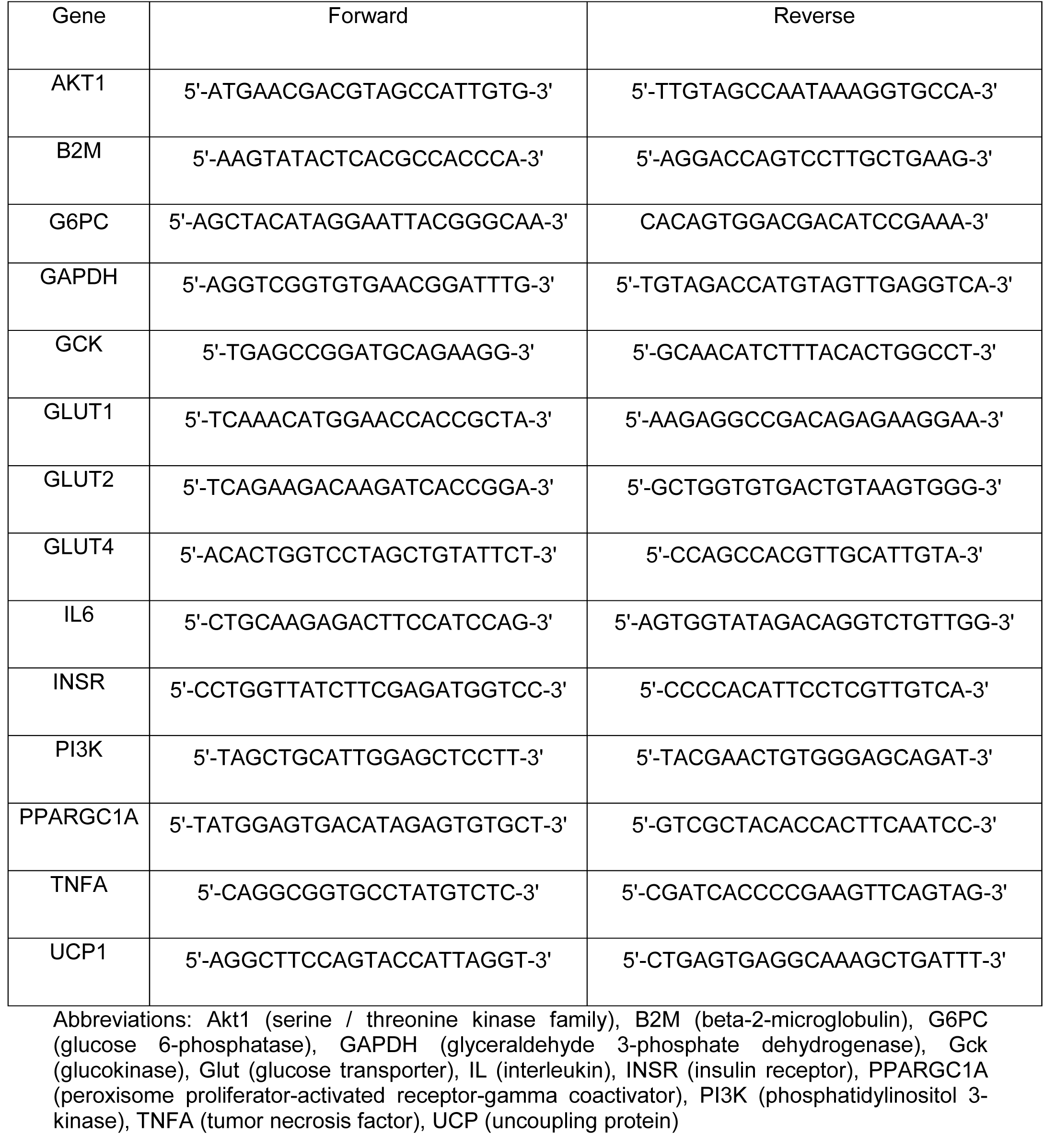
A list of forward and reverse mRNA primers.

### Soluble Aβ_42_ Determination

The hippocampus from one hemisphere was dissected and stored at −80°C until tissue processing. Soluble Aβ_42_ concentrations were determined using the Human / Rat β amyloid ELISA kits (WAKO Chemicals; Cat: 292-64501) according to the manufacturer recommended protocols.

### Statistical Analysis

Prism software Version 10.2 (GraphPad Software, Inc., La Jolla, CA; RRID:SCR_002798) was used for all statistical analyses. A two-way ANOVA was used to test for significance of temperature within a genotype for the ITT, GTT, and MWM learning assays. Temperature treatment differences within the same sex and genotype were determined using a two-tailed Student’s t-test for all remaining assays. Potential outliers were determined with a single Grubb’s test (α=0.05). Data are represented as mean ± SEM and significance was defined as p<0.05.

## Abbreviations

Aβ: amyloid-β
AD: Alzheimer’s disease
AUC: area under the curve
B2M: beta-2- microglobulin
BAFF: B-cell activating factor
FGF21: fibroblast growth factor 21
Tcw: a 42-specific
G6PC: glucose 6-phosphatase
Gck: glucokinase
GLP1: glucagon-like peptide 1
Glut: glucose transporter
GTT: glucose tolerance test
InsR: insulin receptor
ip: intraperitoneal
ITT: insulin tolerance test
MWM: Morris water maze
pg: perigonadal
PGC: peroxisome proliferator-activated receptor-gamma coactivator
PI3K: phosphatidylinositol 3- kinase
Tc: core body temperature
TNF: tumor necrosis factor
UCP: uncoupling protein
VAT: visceral adipose tissue
WAT: white adipose tissue).

## Author Contributions

SAM, MRP, LNS, MFC, EDI, CAF, KQ, and YF assisted with colony maintenance and the experimental assays performed in this manuscript. AB and ERH assisted with experimental design and manuscript revisions. KNH conceived the study, supervised the experiments, analyzed the data, and wrote the manuscript. All authors approved the final version of the manuscript.

## Acknowledgements

We would like to thank Melissa Roberts for conducting the plasma multiplex assay.

## Ethical Statement

Protocols for animal use were approved by the *Institutional Animal Care and Use Committee* at Southern Illinois University School of Medicine (IRB #2022-055) and in accordance with the ARRIVE guidelines.

